# Detergent-Triggered Membrane Remodelling Monitored via Intramembrane Fluorescence De-Quenching

**DOI:** 10.1101/2025.09.24.678259

**Authors:** Claudia M. F. Andrews, Christopher M. Hofmair, Lauryn Roberts, Emily James, Katie Morris, Kevin Kramm, Mark C. Leake, Yue Wang, Steven D. Quinn

**Affiliations:** School of Physics, Engineering and Technology, University of York, Heslington, York, UK. YO10 5DD; PicoQuant, Rudower Chaussee 29, 12849, Berlin, Germany; Department of Biology, University of York, Heslington, York, UK. YO10 5DD; York Biomedical Research Institute, University of York, Heslington, York, UK. YO10 5DD

## Abstract

Detergent-induced membrane solubilization is important for several biotechnological applications including membrane protein isolation, cell lysis and virus inactivation. The thermodynamic details of the underlying process have been previously examined, but the mechanistic details remain largely underexplored owing in part, to a lack of suitable technologies capable of assessing nanoscopic membrane disruption events. Key open questions include: how do detergents remodel the membrane structure at sub-solubilizing concentrations? And what is the sequence of morphological transitions that lead up to solubilization? Here, we introduce a single-colour assay based on the fluorescence de-quenching of membrane-integrated fluorophores as a sensitive and generalizable tool to probe nanoscale membrane remodelling events induced by detergents. We demonstrate, using fluorescence spectroscopy and time-correlated single photon counting, that the widely used detergent Triton X-100 triggers substantial morphological changes at concentrations below its critical micellar concentration. Moreover, by taking advantage of single vesicle fluorescence lifetime imaging and scanning electron microscopy, we reveal that the swelling step involves a morphological transition from spherical vesicles to toroidal structures, providing direct evidence for detergent-driven membrane reorganization prior to solubilization. Our findings support and refine a multistep model of detergent-induced membrane solubilization, positioning fluorescence de-quenching as a tool for detecting conformational intermediates. We show that the fluorescence de-quenching approach performs robustly across multiple cyanine-based probes and experimental conditions and its nanoscale sensitivity provides a platform from which to interrogate membrane perturbations induced by a wide variety of molecular disruptors, including those with important biomedical significance.

## Introduction

Detergent-membrane interactions are critical for a variety of applications including membrane protein extraction and isolation^1^, virus inactivation^2, 3^, cellular drug delivery^4^, forming niosomes in vaccine formulations^5^, and for constructing artificial membranes and supported lipid bilayers^6^. In this context, Triton X-100 (TX-100) is widely used as a gold-standard non-ionic detergent due to its ability to gently solubilize membrane proteins while maintaining their structure and function, making it ideal for biochemical and structural studies^7, 8^. Its relatively low critical micellar concentration (CMC) of ∼0.2-0.3 mM^8, 9, 10^ and mild impact on protein conformation^11^ make it particularly suitable for studies requiring intact protein-lipid interactions, unlike stronger ionic detergents which can disrupt protein functionality. However, the mechanistic details of the TX-100 membrane interaction have remained underexplored, owing in part, to a lack of suitable tools and technologies capable of assessing the interaction. In particular, there remains limited understanding of how detergent molecules initiate structural rearrangements on the nanoscale, what intermediate membrane conformations precede micellization, and how these effects depend on the detergent type, concentration and membrane composition.

In recent years, the emergence of model-membrane systems such as lipid vesicles and supported bilayers^6^ has helped researchers explore the underlying details. Previous biochemical approaches investigating TX-100 interactions demonstrated, for example, that the solubilization rate and effectiveness of the detergent depend on the lipid phase, absolute detergent concentration and the lipid composition^12, 13, 14, 15, 16, 17^. These results, and others, broadly confirm that TX-100 is particularly effective at solubilizing phosphocholine (PC) rich membranes, with solubilization kinetics generally faster in the fluid phase. Gel-phase membranes, in contrast, require higher TX-100 concentrations for complete solubilization, especially as the lipid chain length increases^15^. Additionally, TX-100’s action is inhibited by membrane cholesterol, likely due to the formation of detergent-rich regions that may function as lipid rafts^16, 18^. Molecular dynamics simulations^18, 19, 20^ and phase contrast microscopy experiments^17^ have also provided insights into early-stage permeabilization events, but despite such advancements, challenges remain in dissecting the process at each solubilization stage. Moreover, there is also demand for new tools that can monitor the impact of TX-100 on sub-micron sized membrane systems that are beyond the reach of conventional diffraction-limited optical microscopy approaches.

Nevertheless, based on these experiments a global three-step model has been proposed to describe TX-100-induced membrane solubilization: (1) detergent monomers saturate the membrane, (2) mixed detergent-lipid micelles form leading to membrane fragmentation, and (3) mixed micelles are released into solution^12, 21^. Membrane permeabilization assays based on the influx of calcium into lipid vesicles encapsulating fluorescent indicators have also revealed that transient defects and micropores form on intact vesicles during the process^22^. We also recently showcased dual-colour fluorescence assays based on Förster resonance energy transfer (FRET) between membrane-embedded fluorophores, to support a refined model in which both structural changes and membrane fusion take place prior to step 3^23, 24, 25^. However, FRET-based approaches are not without their limitations. For example, FRET signatures can be challenging to interpret in heterogeneous mixtures as changes in the probe orientation, lipid packing density or local membrane dynamics all influence the Förster distance^26^.

Inspired by these insights, we now introduce a single-colour assay based on the fluorescence de-quenching of membrane-embedded probes as a powerful, but straightforward means to probe and quantify detergent-induced membrane perturbations via enhancements in both fluorescence intensity and probe lifetime. We demonstrate that the de-quenching approach reports on the distance between membrane-embedded fluorophores, and can be implemented using a range of dyes and experimental conditions. We benchmark the de-quenching approach against the existing FRET-based strategy and use it to explore the structural integrity of PC-rich vesicles in response to TX-100. In conjunction with single particle imaging approaches, our observations confirm that TX-100 triggers vesicle swelling and morphological transitions, even at concentrations below the reported CMC, prior to complete solubilization. We expect the fluorescence de-quenching approach to have far-reaching applications in quantitatively reporting on membrane disruption events induced by a wide variety of molecular disruptors including surfactants and proteins with important biomedical significance.

## Methods Materials

1-palmitoyl-2-oleoyl-glycero-3-phosphocholine (POPC) lipids suspended in chloroform, TX- 100, sodium dodecyl sulfate (SDS) and Tween 20 were purchased from Merck. All stock detergent solutions were prepared in 50 mM Tris buffer (pH 8) prior to each use. The lipophilic membrane stains DiI, DiD and DiO were purchased from ThermoFisher Scientific. All lipid and membrane stain stocks were stored at -20°C prior to use. All samples were used directly from the manufacturer without any additional purification.

### Large Unilamellar Vesicle Preparation

POPC vesicles incorporating DiI, DiD or DiO were prepared via the extrusion method. Briefly, the lipids and membrane stains were mixed in chloroform at the levels specified in the main text and the solvent was evaporated under gentle nitrogen flow. The resulting lipid film was then resuspended in 50 mM Tris buffer (pH 8) and vortexed. Unilamellar vesicles were then prepared by passing the solutions at least 21 times through a Mini Extruder (Avanti Research) containing a polycarbonate membrane filter of defined pore size.

### Fluorescence Spectroscopy

Fluorescence emission spectra were acquired under magic angle conditions using a FluoTime 300 (PicoQuant) spectrophotometer. Spectra from vesicles incorporating DiI, DiO and DiD in solution were recorded using excitation wavelengths of 532, 485 nm and 640 nm, respectively. All experiments were performed in 50 mM Tris buffer (pH 8) with a final POPC concentration of 30 μg/mL.

### Time Correlated Single Photon Counting

Time-resolved fluorescence spectroscopy on labelled vesicles in solution was also performed using a FluoTime 300 spectrophotometer equipped with time-correlated single photon counting electronics and a hybrid PMT detector (PMA Hybrid 07, PicoQuant). Time-resolved fluorescence decays were measured under magic angle conditions using pulsed excitation at 532 nm, 485 nm and 640 nm for samples containing DiI, DiO and DiD, respectively. In all cases, repetition rates of 50 MHz were used. Time-resolved fluorescence decays at the maximal intensity emission wavelengths were collected until 10^4^ photon counts accumulated at the decay maximum. Fluorescence decay curves were then fitted by iterative reconvolution of the instrument response function and the observed fluorescence decay using a bi-exponential decay function of the form I_t_ = ae^-t/*τ*1^+ be^-t/*τ*2^, where I_t_ is the intensity at time, t, normalized to the intensity at t = 0, *τ*_1_ and *τ*_2_ represent the fluorescence lifetimes of fast and slow decay components, and a and b are the associated fractional amplitudes. We note that bi-exponential fits were applied based on the convergence of the reduced chi-squared to the experimental data. All experiments were performed in 50 mM Tris buffer (pH 8) with a final POPC concentration of 30 μg/mL. The variation in amplitude weighted average lifetimes was fitted to a Hill model of the form 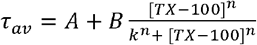, where A and B are the measured lifetimes at the start and end of the titration, k is the half-maximal concentration constant and n is the Hill coefficient.

### Single Vesicle Fluorescence Lifetime Imaging (svFLIM)

svFLIM measurements were performed using a Luminosa single photon counting confocal microscope (PicoQuant) using the dedicated FLIM workflow available in the system. Briefly, 50 μL droplets of vesicles in solution were pipetted onto high precision cover glasses coated in 1 % poly-l-lysine. Excitation was provided by a pulsed diode laser emitting at 532 nm with pulse width of 76 ps and repetition rate of 20 MHz (LDH-D-FA-530L) at an output power of 0.5 µW (measured after the main dichroic mirror). Images were acquired with a FLIMbee galvo scanner. The emission was collected with a 60x 1.20 numerical aperture objective lens (Evident). A single-photon avalanche diode (SPAD; Excelitas AQRH-14) and suitable optical filters were used to detect fluorescence photons in the spectral range between 545 and 615 nm. Photon detection events were time-correlated to excitation pulses using a time-correlated single-photon counting device (MultiHarp 150 8P, PicoQuant). Fluorescence decays were fitted to bi-exponential decays using the Luminosa software (PicoQuant).

### Scanning Electron Microscopy (SEM)

SEM was performed using a JEOL JSM 7800-F system operating at 5 kV. Vesicles were prepared in 50 mM Tris (pH 8) containing TX-100 at the specified concentrations, diluted in deionized water, and vortexed prior to imaging. 2 μL volumes of the vesicle solution were then added to a clean silicon substrate and the solution evaporated. Reference samples without TX-100 were prepared under the same conditions. All substrates were then sputtered with an 8 nm copper layer for charge dissipation purposes before loading into the microscope. Vesicle diameters were determined using ImageJ, where automated analysis of black and-white binary images enabled separation of regions of white pixels against a dark background. Vesicle circularity was measured via 4p(A/p^2^), where A is the observed area and p is the perimeter.

## Results and Discussion

### Fluorescence de-quenching as a probe of TX-100 induced vesicle solubilization

Large unilamellar vesicles composed of 99 % POPC lipids and 1 % of the lipophilic cyanine derivative DiI were first prepared as outlined in the Methods, and are schematically illustrated in **Figure 1a**. As previously reported, POPC is abundantly found in mammalian membranes and was used here to provide a synthetic mimetic^27^. Here, a mean DiI-DiI separation distance of < 2 nm was achieved, leading to a high level of fluorescence self-quenching. We hypothesised that detergent-induced structural rearrangements, such as swelling, fragmentation and lysis, would lead to nanoscale increments in the average dye-dye separation distance that, in turn, could trigger fluorescence de-quenching and quantifiable changes to the mean DiI fluorescence emission intensity and lifetime as illustrated in **Figure 1a**.

**Figure 1.**
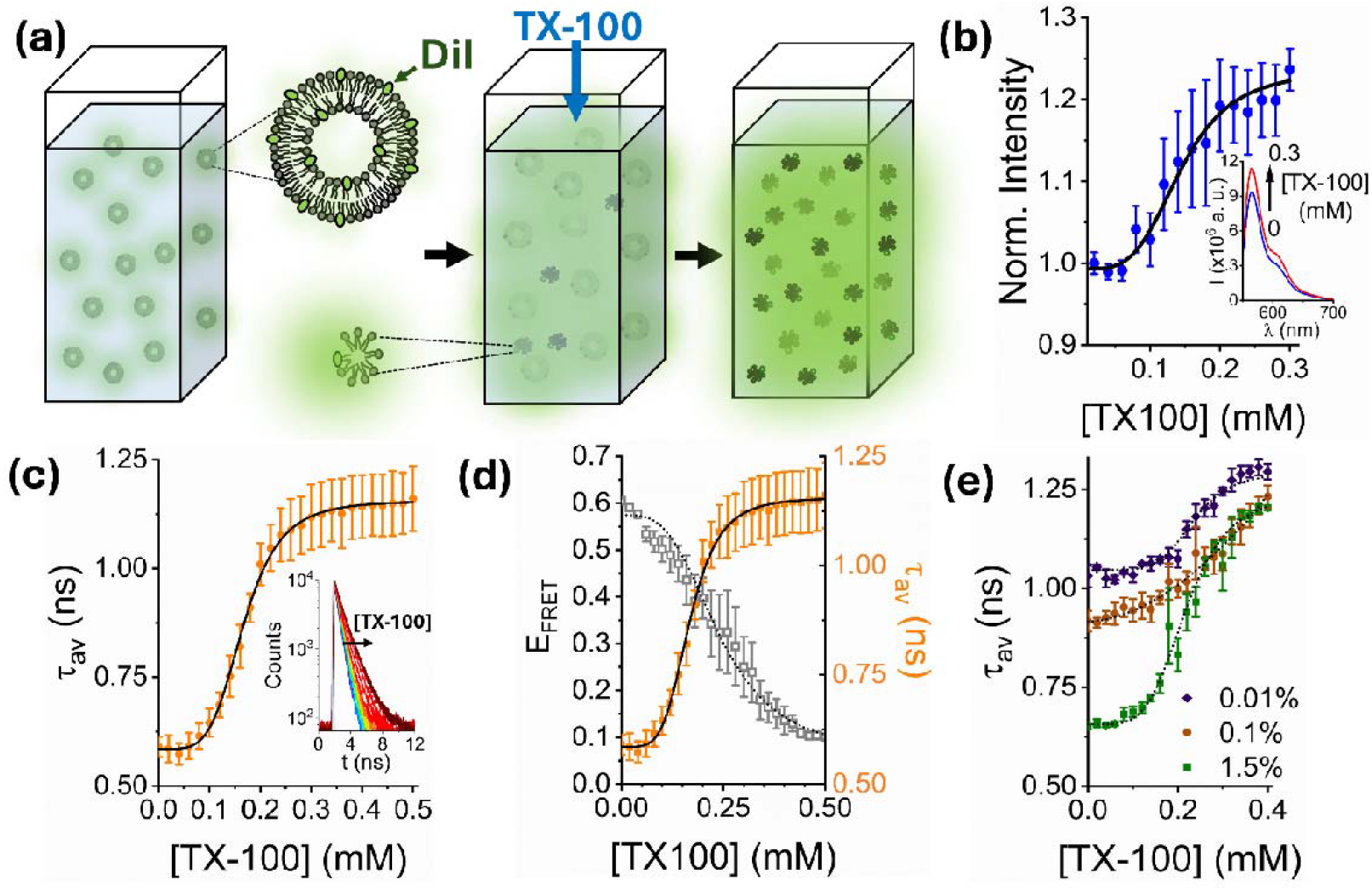
TX-100 induced vesicle solubilization monitored by fluorescence de-quenching. (a) Schematic illustration of the assay. Injection of TX-100 into solution containing DiI-labelled POPC vesicles induces vesicle solubilization, triggering an increase in the relative DiI-DiI distance and de-quenching. (b) Variation in the integrated fluorescence emission intensity of 200 nm sized vesicles containing 1 % DiI in the absence and presence of TX-100. Inset: representative variation in fluorescence emission spectra in the absence and presence of 0.3 mM TX-100. The solid black line represents a Hill model fit (χ^2^ = 0.92) with a half maximal concentration constant, k, of 0.14 ± 0.03 mM. (c) The corresponding variation in *τ*_av_ across the titration with corresponding Hill fit (solid black line; χ^2^ = 0.99; k = 0.17 ± 0.01 mM). Inset: the corresponding fluorescence decays and instrumental response function (grey). (d) Variation in *τ*_av_ associated with vesicles containing 1 % DiI as a function of TX-100 concentration compared to the variation in E_FRET_ obtained from vesicles containing 0.1 % DiI and 0.1 % DiD under identical conditions. The dashed line represents the corresponding Hill model (χ^2^ = 0.99; k = 0.24 ± 0.01 mM). (e) Variation in *τ*_av_ and Hill fits (dashed lines) associated with vesicles containing 0.01 % (χ^2^ = 0.98; k = 0.23 ± 0.01 mM), 0.1 % (χ^2^ = 0.99; k = 0.21 ± 0.01 mM), and 1.5 % DiI (χ^2^ = 0.99; k = 0.22 ± 0.01 mM) as a function of TX-100. In all cases, data points represent the mean values from three separate experimental runs and error bars denote the standard error of the mean.

200 nm-sized vesicles incorporating DiI were prepared via extrusion, and their steady-state fluorescence emission spectra were recorded as a first step to characterize their interaction with TX-100 above and below the reported CMC. As the concentration of TX-100 was progressively increased, a 24 ± 2 % increase in the integrated DiI emission intensity was observed across the titration, consistent with a de-quenching mechanism (**Figure 1b**). To confirm a de-quenching process, we also evaluated the amplitude-weighted average lifetime, *τ*_av_, of membrane-bound DiI using time-correlated single photon counting. Here, *τ*_av_ progressively increased with TX-100 concentration consistent with a progressive de-quenching of DiI (**Figure 1c**). In all cases, and in line with previous observations^23^, the time-resolved fluorescence decays were best fit to a bi-exponential decay model after re-convolution with the instrument response function, representing variations in the dye’s local environment^22^. In the absence of TX-100 we recorded a *τ*_av_ = 0.59 ± 0.04 (± SD) ns, representative of quenched DiI. A 2-fold increase in *τ*_av_ was then observed as TX-100 was titrated above and below the CMC (**Figure 1c**). Variations in *τ*_av_ as a function of TX-100 concentration fitted well to a Hill model, revealing a half-maximal concentration of 0.17 ± 0.01 mM, which we note is comparable in magnitude to the reported CMC. Upon further inspection, we found that both fast and slow lifetime components increased across the titration, though their relative amplitude weighted percentage contributions remained largely invariant (**Figures S1 and S2**). The concentration-dependent increase in *τ*_av_ showed a similar half-maximal concentration to that obtained from the solubilization of comparably sized vesicles labelled with 0.1 % DiI and 0.1 % DiD evaluated using the previously-demonstrated FRET-based sensing approach^22, 23, 24^ (**Figure 1d**). Here, a TX-100 induced distance increase between donor (DiI) and acceptor (DiD) probes leads to a progressive reduction in the apparent FRET efficiency, E_FRET_, across the titration. The anti-correlation observed between the *τ*_av_ signal obtained from the single-colour assay and the apparent FRET efficiency from the dual-colour approach, provides confidence that the de-quenching approach quantitatively reports on perturbations on the nanoscale. Unlike the FRET-based approach, however, the de-quenching assay avoids donor-acceptor stoichiometry dependence and spectral bleed-through, enabling a direct readout of membrane disruption. We also note that the amount of DiI per vesicle was optimized (1 %) to maximize the magnitude of the de-quenching response across the interaction. In this context, the initial amplitude-weighted lifetime progressively increased towards a de-quenched state as the dye concentration was reduced to 0.1 % and 0.01 %, respectively (**Figure 1e**). The initial lifetime values were, however, comparable when the vesicles contained 1 % and 1.5 % DiI. In all cases, and in line with the data shown in **Figure 1c**, a progressive increase in the probe lifetime was observed, even at concentrations below the reported CMC, and a final end-point (1.2-1.3 ns) was reached.

### Fluorescence de-quenching as a generalizable assay

To assess whether the fluorescence de-quenching approach was applicable beyond the DiI system, we also evaluated the response of 200 nm sized vesicles containing DiO and DiD. Like DiI, DiO and DiD are long-chain dialkylcarbocyanines with emission excitation/emission maxima at 484/501 nm and 644/665 nm, respectively^28^. As the concentration of TX-100 was progressively increased, the integrated emission intensity of vesicles containing 1 % of each dye also progressively increased, in line with our previous observations (**Figure 2a, b**). In both cases, the fluorescence decay curves also shifted to longer decay times as evidenced by a progressive increase in the *τ*_av_ signatures (**Figure 2c, d**). Here, the initially quenched lifetimes of DiO (0.51 ± 0.01 ns) and DiD (0.97 ± 0.01 ns) increased to 0.79 ± 0.01 ns and 1.87 ± 0.03 ns respectively as TX-100 was injected above and below the CMC. In both cases the half-maximal concentration requirements to achieve solubilization were identical (k = 0.19 ± 0.01 mM) and in agreement with our observations from DiI. We note that the magnitude of the lifetime changes between the start- and end-point of the titration increased in the order DiO > DiI > DiD (**Figure 2e**), which we speculate might reflect differences in the dye-lipid interaction strength, the extent of dye interactions within the membrane, and/or variations in the degree of dye insertion into the lipid bilayer.

**Figure 2.**
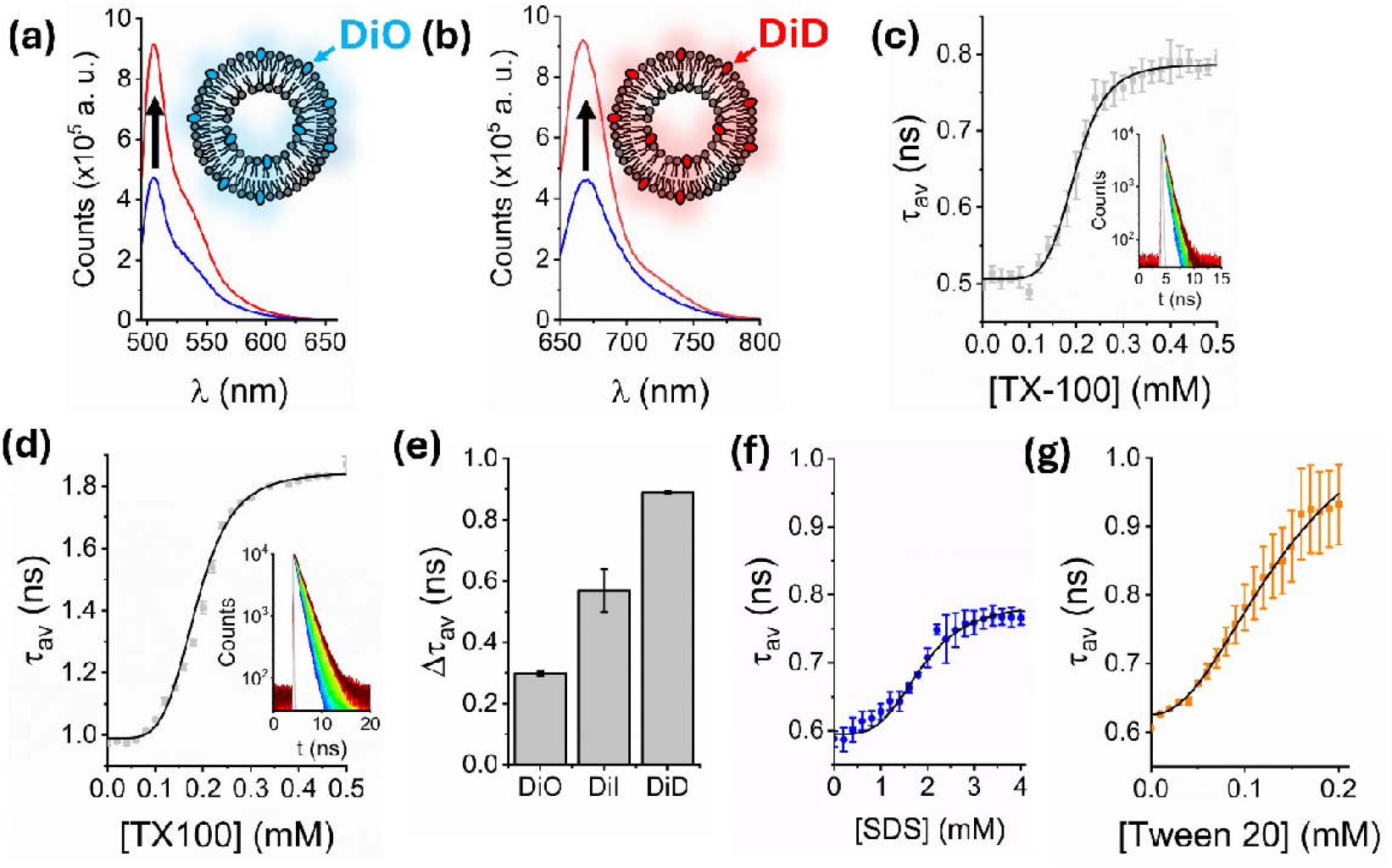
Expanding the scope of fluorescence de-quenching. Variation in the integrated fluorescence emission intensity of 200 nm sized POPC vesicles containing (a) 1 % DiO and (b) 1 % DiD in the absence (blue) and presence (red) of 0.3 mM TX-100. Insets: schematic illustrations of the vesicles. Representative variation in *τ*_av_ as TX-100 was progressively added to vesicles containing (c) 1 % DiO and (d) 1 % DiD. Insets: the corresponding time-resolved fluorescence decays. The solid black lines represent Hill fits to the experimental data (χ^2^ = 0.99; k = 0.19 ± 0.01 mM in both cases) and the solid grey lines indicate the instrumental response functions. (e) Comparative bar plot summarizing the relative magnitude of the de-quenching signal between the start and end of the titration as a function of dye at 1 %. (f) Representative variation in *τ*_av_ obtained from POPC vesicles labelled with 1 % DiI in the absence and presence of SDS with Hill fit shown in the solid black line (χ^2^ = 0.96; k = 1.9 ± 0.01 mM). (g) The corresponding *τ*_av_ data and fit (χ^2^ = 0.96; k = 0.14 ± 0.09 mM) obtained from vesicles containing 1 % DiI in the presence of Tween 20. Error bars represent the standard error of the mean from three separate experimental runs.

Having established that the de-quenching assay operates across a range of fluorescent probes, we next evaluated the response of DiI loaded POPC vesicles to other surfactants including sodium dodecyl sulfate (SDS) and Tween 20 to gauge its utility across a range of molecular disruptors. Unlike TX-100, SDS is an anionic detergent with a linear alkyl chain, a sulfate head group, and CMC of ∼6-8 mM, making it highly denaturing^24^. Tween 20, like TX-100, is also non-ionic, with a polyoxyethylene sorbitan ester structure, a CMC of ∼0.06 mM, and is generally considered mild upon comparison with SDS^22^. Under both conditions, the initial amplitude-weighted average lifetimes were comparable to those previously reported, indicating reproducibility of the starting material. Upon addition of SDS, we observed a progressive increase in *τ*_av_ from 0.59 ± 0.01 to 0.77 ± 0.01 ns, with a half-maximal concentration constant of 1.9 mM (**Figure 1f**). By contrast, the lifetime increased to 0.93 ± 0.01 ns upon addition of only 0.2 mM Tween 20 (**Figure 1g**). The relative difference in the magnitude of the fluorescence lifetime shift between the two detergents could arise because of distinct interactions with the vesicle membrane and/or due to variations in the final micellar forms. For example, SDS is an anionic surfactant that likely disrupts membrane packing to a lesser extent than the non-ionic surfactant Tween 20, which may integrate more effectively into the bilayer and alter the local environment of DiI. SDS and Tween 20 also form micelles with distinct structural characteristics; SDS generally induces small, spherical mixed detergent-lipid micelles which have a high negative surface charge and tight packing, whereas Tween 20 induced micelles are larger and less uniformly shaped with the detergent arranged in a more loosely packed environment^29, 30^. Nevertheless, in all cases it is striking to note that the de-quenching signal occurs at concentrations below the respective detergent’s critical micellar concentration, indicating that the assay is sensitive to conformational changes such as vesicle swelling which occurs before solubilization and release of mixed detergent-lipid micelles.

### Triton X-100 triggers vesicle swelling and morphological transitions

To assess whether the observed variations in fluorescence intensity and lifetime could be assigned to vesicle swelling, we performed an array of single-vesicle imaging experiments. We first performed fluorescence lifetime imaging (FLIM) to assess the lifetime distributions from single DiI-labelled vesicles. Here, 200 nm sized POPC vesicles containing DiI at 1 % were non-specifically attached to a poly-l-lysine-coated coverslip and the variation in lifetime distribution across the surface, and across single vesicles, was monitored before and after the addition of TX-100 at sub-solubilizing concentrations (**Figure 3a**). It is important to note that the immobilized vesicles were also spatially isolated on the surface, minimizing the possibility of fusion. In the absence of TX-100, we observed ∼200-300 fluorescent foci per 190 × 190 μm^2^ field of view, representing individual non-specifically bound vesicles (**Figure 3b**). In line with our prior ensemble-based measurements, the distribution of lifetime values from hundreds of surface-immobilized vesicles also fitted well to a bi-exponential model comprising fast (0.76 ± 0.16 ns) and slow (2.01 ± 0.14 ns) components and an amplitude-weighted average lifetime of 0.99 ± 0.3 ns (**Figure 3c**). As 0.15 mM TX-100 was added, we observed a substantial increase in both lifetime components (*τ*_1_ = 1.32 ± 0.09 ns and *τ*_2_ = 2.27 ± 0.04 ns), and *τ*_av_ increased by ∼68 % to 1.67 ± 0.34 ns (**Figure 3c**) while the total number of fluorescent foci per field of view, and therefore the number of vesicles on the surface, remained largely invariant (**Figure S3**). The lifetime increase observed across single vesicles at low TX-100 concentrations could not therefore be attributed to complete solubilization of the vesicles or lipid loss, but rather structural changes taking place within the intact vesicles. We assigned the de-quenching signal observed here to detergent-induced remodelling of the membrane that involves vesicle swelling and/or morphological transitions with minimal loss of lipid material to the bulk solution. We note that the distribution of individual lifetime components obtained across single vesicles was generally Gaussian distributed with a full-width at half-maximum of ∼0.2 ns, indicating a lack of fused, aggregated or significantly perturbed species on the surface (**Figure 3d, e**). Our observations were also reflected when random sampling of the mean component lifetimes per vesicle revealed a 1.5-fold increase in *τ*_1_ (**Figure 3f**) and a 1.3-fold increase in *τ*_2_ (**Figure 3g**). Furthermore, the corresponding ratio of amplitudes associated with the fast and slow components (A(*τ*_1_)/A(*τ*_1_)) decreased from 4.4 ± 1.1 to 2.4 ± 0.8 (**Figure 3h**), indicating that TX-100 alters the lipid microenvironment, disrupting dye-dye interactions that favour the fast lifetime component.

**Figure 3.**
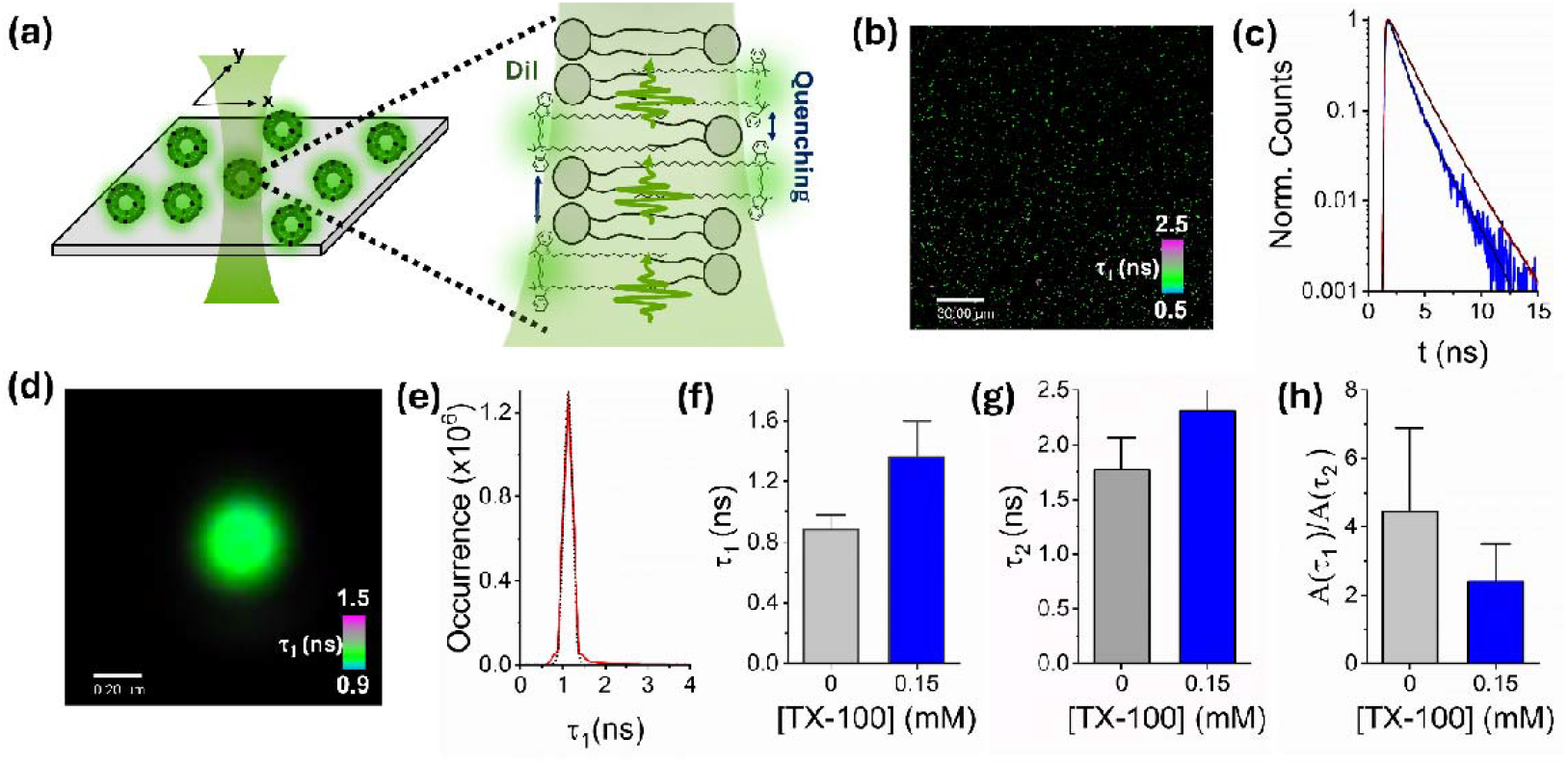
TX-100 induces the structural remodelling of single immobilized vesicles. (a) Schematic illustration of the FLIM approach. Individual vesicles incorporating DiI are immobilized and spatially isolated on a surface, where they are excited by pulsed excitation in a confocal geometry. (b) Representative FLIM image of surface-immobilized POPC vesicles incorporating 1 % DiI showing variations in fast component, *τ*_1_ across the field of view. (c) Variation in TCSPC FLIM decays recorded from vesicles in the absence (blue) and presence of 0.15 mM TX-100 (red). Solid black lines represent bi-exponential fits after reconvolution with the instrument response function. (d) Representative FLIM image of a single DiI-vesicle (no detergent) showing variations in fast component, *τ*_1_ across the structure. (e) The corresponding distribution of *τ*_1_ values across the vesicle surface. The dotted black line corresponds to a Gaussian fit with centre of 1.12 ns, full width at half maximum of 0.2 ns and χ^2^ = 0.99. Comparative bar charts summarizing the mean variations in (f) *τ*_1_, (g) *τ*_2_ and (h) A(*τ*_1_)/A(*τ*_2_) obtained from N = 5 randomly selected immobilized vesicles. Error bars correspond to the standard deviations.

To confirm the detergent-induced swelling of intact vesicles, and to assess whether or not morphological transitions were taking place, we also probed vesicle sizes and structures in response to TX-100 via scanning electron microscopy. Here, we assessed the size distribution of large unilamellar vesicles in the absence and presence of TX-100. SEM micrographs revealed that freshly prepared vesicles were predominantly spherical (mean circularity = 0.73; standard deviation = 0.28, standard error of the mean = 0.04) (**Figure 4a, b, c**) with a mean diameter of 750 nm (standard deviation = 275 nm, standard error of the mean = 40 nm, N = 50) (**Figure 4d**). After treatment with 0.3 mM TX-100, the vesicles were also spherical (mean circularity = 0.84; standard deviation = 0.09, standard error of the mean = 0.02) (**Figure 4d, e**) but the size distribution broadened, and the mean diameter was centred on 1173 nm (standard deviation = 379 nm, standard error of the mean = 54 nm, N = 50), representing a 56 % increase. At the 0.05 level, an unpaired sample t test indicated that the difference between population means is significantly different. We also note that exposure to TX-100 induced a morphological transition from intact spherical vesicles to toroidal-like structures (**Figure 4e-h**). At sub-solubilizing concentrations, we expect TX-100 to insert into the membrane bilayer, disturbing lipid packing and increasing fluidity without fully disrupting the intact vesicle. This partial solubilization then likely leads to the formation of intact donut-like toroidal structures as the membrane reorganizes to minimize edge energy while accommodating detergent-lipid micellar phases. In short, we hypothesise that this transformation reflects a transition between bilayer and mixed-micelle structures, driven by TX-100’s ability to destabilize the bilayer architecture. We note that the SEM sample preparation involved using a thin (8 nm) conductive layer, but as previous studies indicate, this minimally alters the particle morphology^31^. Taken in conjunction with our single-particle FLIM data, the presented work is broadly supportive of a solubilisation model that encompasses TX-100 induced vesicle swelling and morphological transitions, prior to complete micellization.

**Figure 4.**
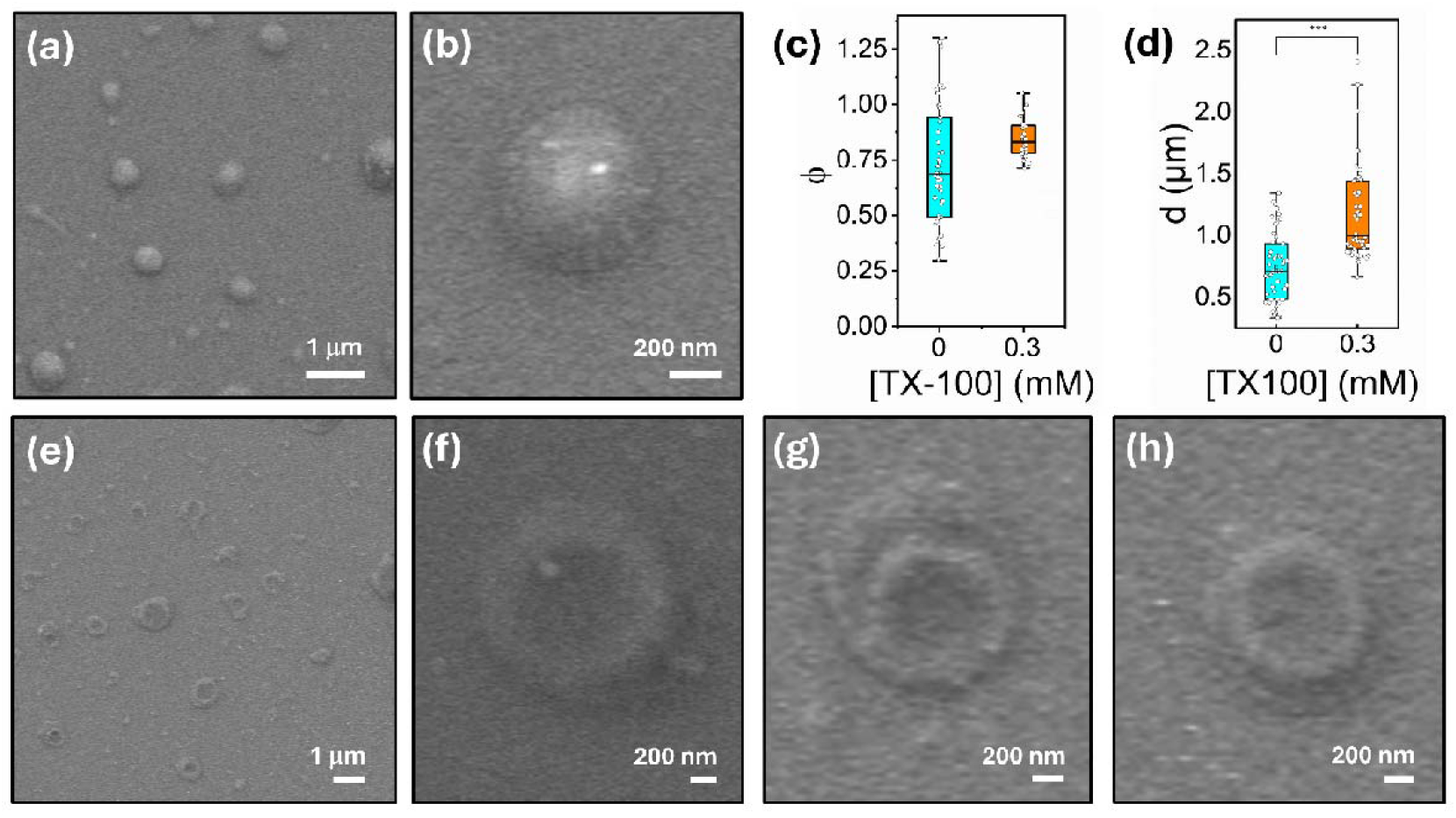
TX-100 induces vesicle swelling and morphological transitions. (a,b) Representative SEM images of freshly prepared vesicles in the absence of TX-100. Comparative bar plots summarizing the variation in (c) circularity and (d) particle diameter for vesicles in the absence and presence of 0.3 mM TX-100 (N = 50). ***p<0.05 as determined by an unpaired sample t test. (e-h) Representative SEM images of freshly prepared vesicles in the presence of 0.3 mM TX-100.

A remarkable outcome of this study is the demonstration that fluorescence de-quenching can sensitively report on detergent-induced conformational expansion and morphological restructuring of individual intact vesicles in solution. Furthermore, these effects were observed at concentrations approaching the CMC, highlighting the capacity of nonionic detergents to induce significant structural rearrangements in highly curved vesicles. While the current data does not report on the initial dynamics of single TX-100 molecules directly interacting with the membrane, and further work in this area is highly desirable to quantify the precise mechanistic details of the initial interaction, our findings broadly support a model of TX-100 induced solubilization that involves structural remodelling of the intact vesicle prior to lysis. In this context, fluorescence de-quenching is a powerful single-colour alternative to dual-colour FRET-based approaches, and opens a platform for enabling dynamic, nm-scale insights into membrane behaviour without requirements for complex fluorophore pairing. Further advantages of the technique include its ability to operate across a range of fluorophores and surfactant conditions, and to enable interrogation of vesicle structure on a vesicle-by-vesicle basis, bypassing ensemble averaging which often obscures heterogeneous or transient events within vesicle populations.

Our data reveals that TX-100 induces vesicle swelling and the formation of non-spherical, toroidal-like structures prior to membrane rupture and the formation of mixed detergent-micelles. This multistep solubilization mechanism is consistent with previous observations in giant unilamellar vesicles (GUVs), where TX-100 was shown to induce long-lived pores and substantial vesicle shape changes^32, 33, 34, 35^. Indeed, a direct comparison between the data obtained from large unilamellar vesicles and giant vesicles composed of POPC revealed common features after incubation with TX-100, including similar concentration requirements to achieve solubilization, morphological changes below the detergent’s CMC, and evidence of the formation of toroidal-like structures (**Figure S4**). Both systems also displayed quenched initial lifetimes, bi-exponential decay behaviour and an increase in average lifetime under matched detergent conditions. Interestingly, we identified that the half-maximal concentration for solubilization in the case of the giant vesicles was 0.18 ± 0.01 mM, which we note is in line with the value obtained from the vesicles of 200 nm diameter. In both scenarios, the high curvature of the vesicles likely destabilizes upon detergent intercalation, leading to a reduction in membrane tension that facilitates global deformability and morphological transitions. This scenario aligns with theoretical models in which TX-100 – due to its amphipathic, flat wedge-like geometry and near-zero spontaneous curvature - intercalates symmetrically into the bilayer. Through flip-flop, lipid packing may then be compromised, membrane asymmetry disrupted and conditions become favourable for non spherical morphologies and, subsequently, mixed micelle formation. Irrespective, the fluorescence de-quenching approach on highly curved, sub micron sized vesicles, is complementary to previous optical microscopy experiments involving giant unilamellar vesicles, and the combined data suggests a common general mechanism of solubilization.

Our findings also corroborate previous work that demonstrates that TX-100 perturbs local curvature and induces the formation of worm-like or tubular vesicles, invaginated forms, and other non-spherical intermediates^33, 34, 36, 37^. Similarly, the observed structural transformations also echo earlier reports of TX-100–induced swelling and shape changes in chloroplast membranes and human red blood cells^38, 39^, reinforcing the broader relevance of our observations. An important mechanistic implication of our study, however, is the observation that significant membrane deformations occur at lipid-to-detergent ratios of ∼2,000:1— equivalent to < 100 detergent monomers per vesicle. This supports the hypothesis that insertion and accumulation of individual TX-100 molecules, even at low concentrations below the CMC, can initiate conformational transitions.

It is also worth re-emphasizing the key advantages of the fluorescence de-quenching approach. First, individual fluorescently tagged vesicles can be interrogated on a vesicle-by-vesicle basis thereby bypassing major limitations associated with ensemble averaging tools. Second, the assay operates well on the nanoscale, providing sensitivity comparable to FRET based measurements, but without the complexities associated with spectral overlap and bleed-through. Indeed, we have established that the combination of ensemble and single-vesicle spectroscopy approaches based on fluorescence de-quenching can be used to reveal and monitor precise molecular level events that underpin detergent-induced vesicle solubilization in vitro, and we identify that TX-100 alters the structure of both freely diffusing and surface-immobilized vesicles via a mechanism comprising swelling and the formation of toroidal-like structures prior to complete solubilization. Our observations provide new mechanistic insight for how solubilizing detergents perturb and damage highly curved vesicles and may be directly relevant to biotechnological applications where conformational control and manipulation of the membrane is vital. We also expect the presented approach to find general utility for unveiling vesicle structural changes in response to perturbative agents, including additional surfactants, disruptive proteins and antiviral agents beyond the test cases highlighted here.

## Conclusions

The combination of ensemble and single-vesicle fluorescence de-quenching approaches provides a powerful platform to study detergent-induced membrane remodelling. The ability to quantify structural changes in vesicles in response to detergents at both the population and individual levels offers utility for studying the role of perturbative agents more broadly. Our findings offer mechanistic insight into how TX-100 acts on highly curved vesicles, revealing that the detergent triggers vesicle swelling and transitions from intact spherical to toroidal morphologies, before complete solubilization. This now opens potential implications for biotechnological applications where controlled membrane remodelling is desirable. We anticipate the toolbox presented here will also be valuable in assessing vesicle responses to other classes of molecular disruptors, including membrane-active peptides, surfactants, pharmacological agents and proteins with important biomedical significance.

## Acknowledgements

The authors acknowledge the Engineering and Physical Sciences Research Council (EP/V034030/1 and EP/V047663/1) and Alzheimer’s Research UK (RF2019-A-001) for support. We also thank Dr. Lara Dresser (University of York, UK) for help with initial exploratory work, Laser 2000 (Joshua Seagrave) and Picoquant GmbH for use of the Luminosa equipment and the Bioscience Technology Facility at the University of York.

## Declarations

Co-author K. K. is an employee of PicoQuant.

## Supporting Information

Time-resolved fluorescence data, representative FLIM and SEM images.

