## Supporting Information for "Detergent-Triggered Membrane Remodelling Monitored via Intramembrane Fluorescence De-Quenching"

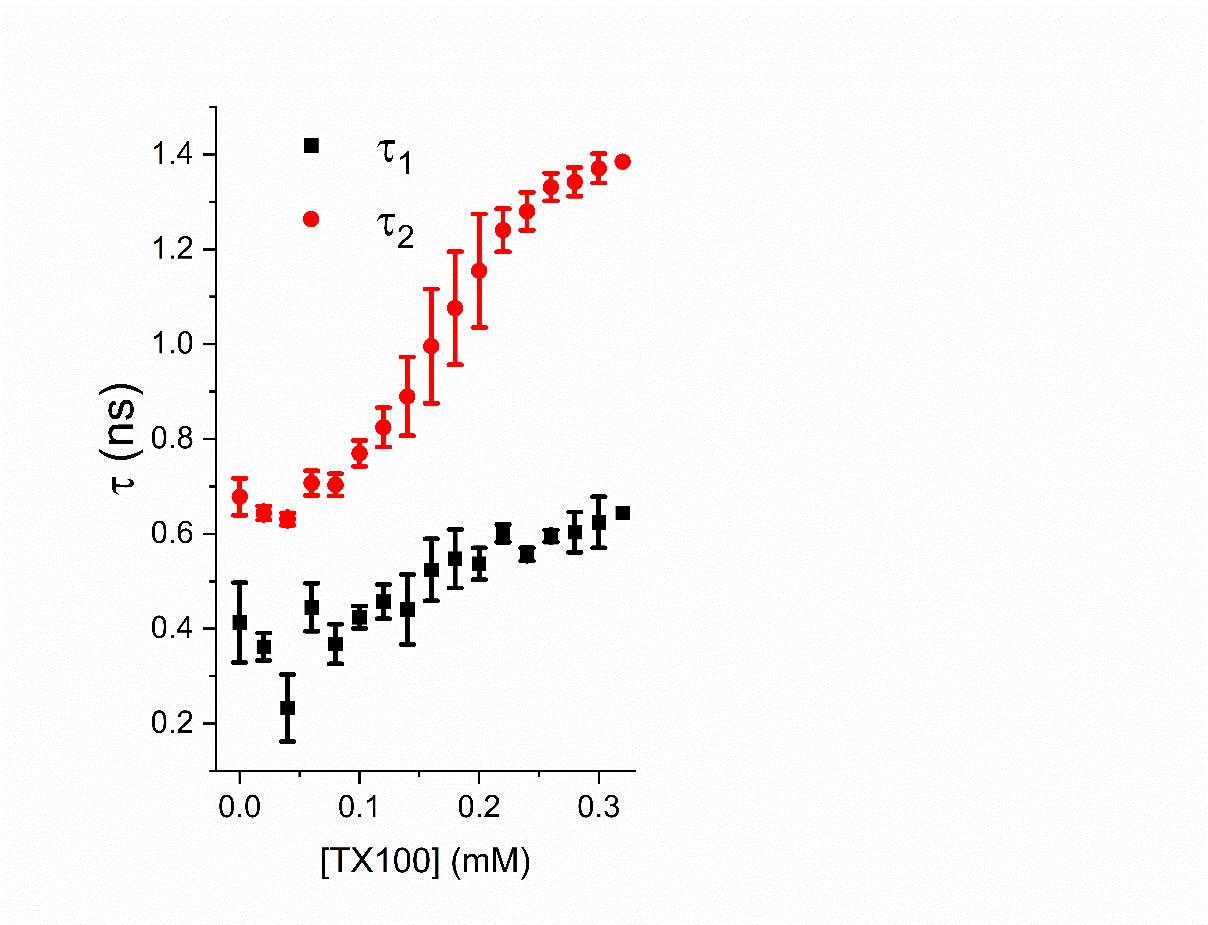

**Figure S1**. Representative variation in fast (τ_1_)and slow (τ_2_) lifetime components obtained from POPC vesicles containing 1 % DiI across the titration. Data points represent the mean and standard error of the mean from three separated experimental runs. Solution conditions: 50 mM Tris, pH 8.

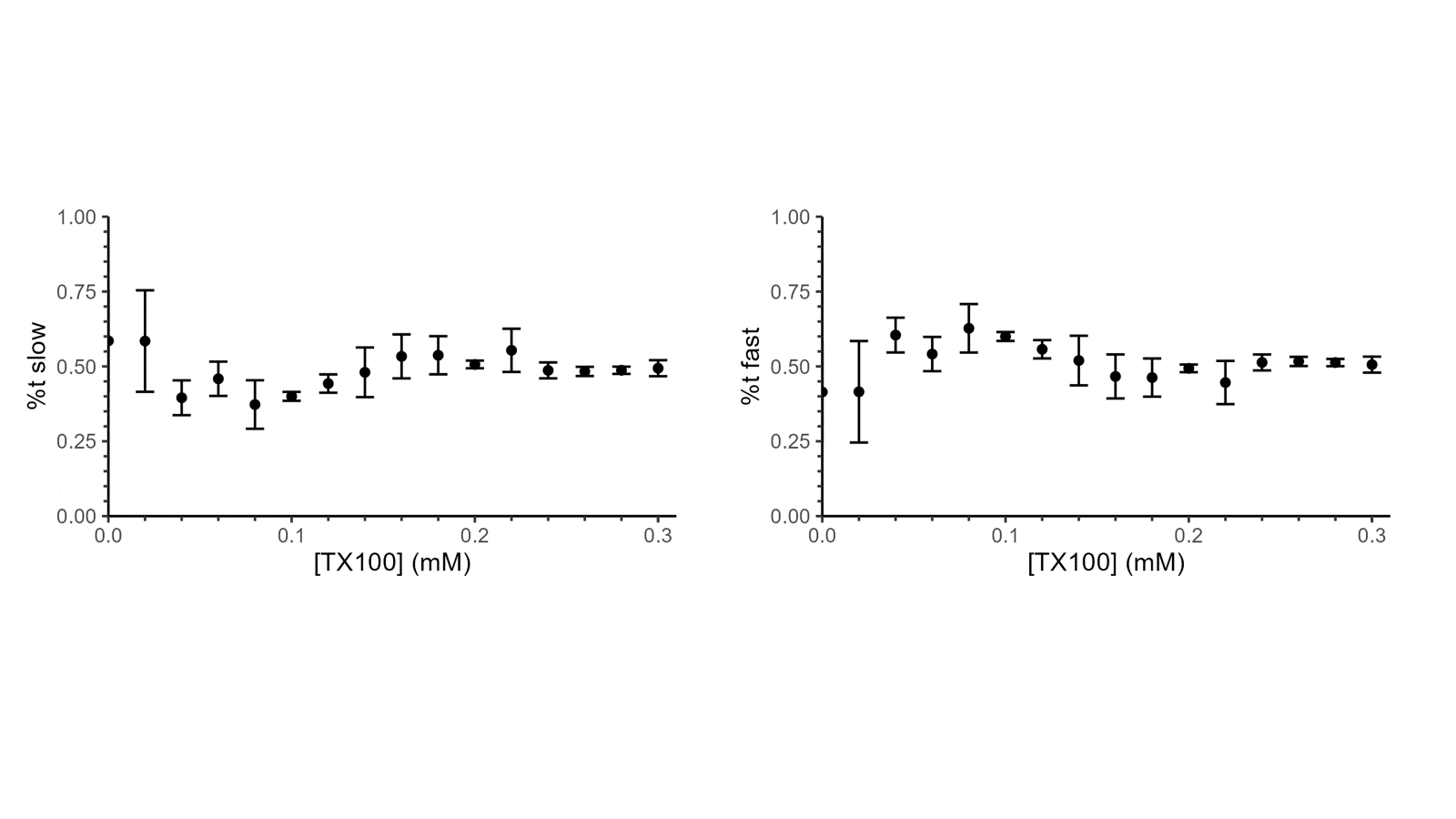

**Figure S2**. Representative variation in % contributions of the fast and slow lifetime components obtained from POPC vesicles containing 1 % DiI across the titration. Data points represent the mean and standard error of the mean from three separated experimental runs. Solution conditions: 50 mM Tris, pH 8.

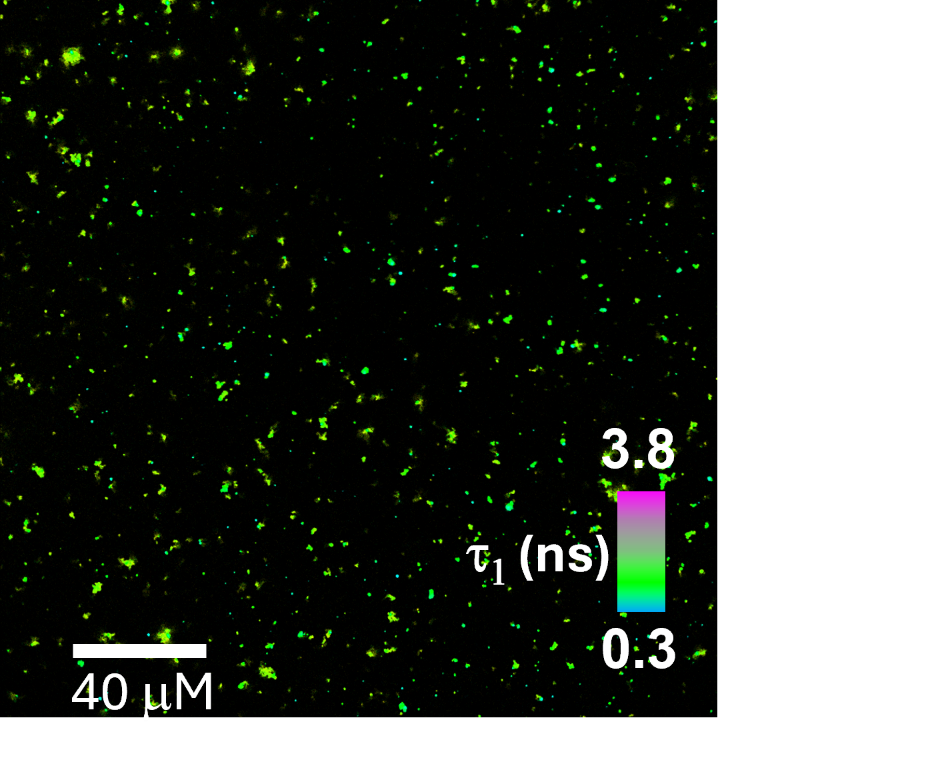

**Figure S3.** Representative FLIM image of surface-immobilized POPC vesicles incorporating 1 % DiI after addition of 0.15 mM TX-100 showing variations in fast component, τ_1_ across the field of view.

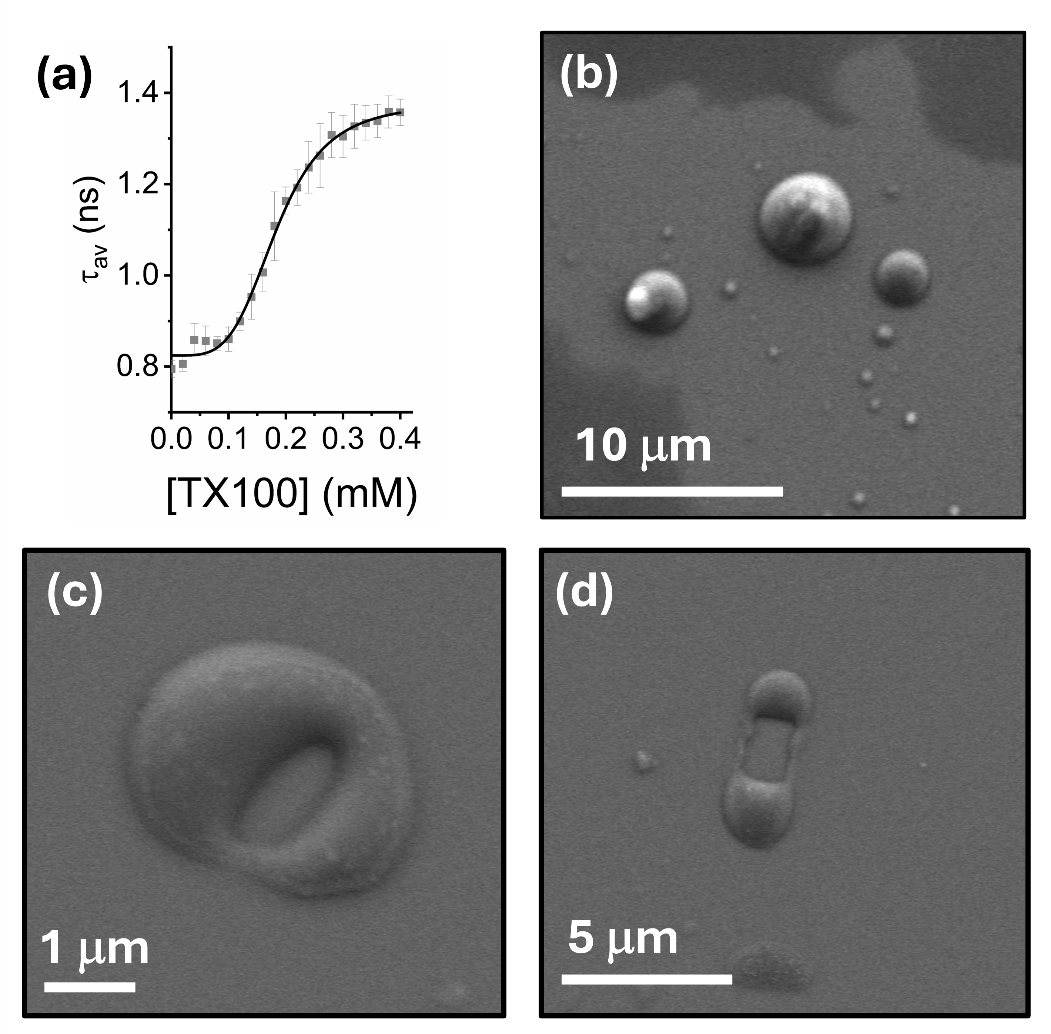

**Figure S4. TX-100 induces vesicle swelling and morphological alterations in giant unilamellar vesicles.** (a) Representative variation in τ_av_ as TX-100 was progressively added to freshly prepared giant unilamellar vesicles (GUVs) containing 1 % DiI. The solid black line represents a Hill fit to the experimental data (χ^2^ = 0.99; k = 0.18 ± 0.01 mM). (b) Representative SEM images of freshly prepared GUVs in the absence and (c,d) presence of 0.3 mM TX-100.
